# Dissociation between Frontal and Temporal-Parietal Contributions to Connected Speech in Acute Stroke

**DOI:** 10.1101/824300

**Authors:** Junhua Ding, Randi Martin, A. Cris Hamilton, Tatiana T. Schnur

**Author notes:** Corresponding author at: Department of Neurosurgery, Baylor College of Medicine, One Baylor Plaza, Houston, Texas 77030, United States. Please cite as: Ding, J., Martin, R., Hamilton, A.C., & Schnur, T.T. (in press). Dissociation between frontal and temporal-parietal contributions to connected speech in acute stroke. Brain.

## Abstract

Humans are uniquely able to retrieve and combine words into syntactic structure to produce connected speech. Previous identification of focal brain regions necessary for production focused primarily on associations with the content produced by speakers with chronic stroke, where function may have shifted to other regions after reorganization occurred. Here, we relate patterns of brain damage with deficits to the content and structure of spontaneous connected speech in 52 speakers during the acute stage of a left hemisphere stroke. Multivariate lesion behavior mapping demonstrated that damage to temporal-parietal regions impacted the ability to retrieve words and produce them within increasingly complex combinations. Damage primarily to inferior frontal cortex affected the production of syntactically accurate structure. In contrast to previous work, functional-anatomical dissociations did not depend on lesion size likely because acute lesions were smaller than typically found in chronic stroke. These results are consistent with predictions from theoretical models based primarily on evidence from language comprehension and highlight the importance of investigating individual differences in brain-language relationships in speakers with acute stroke.

## Introduction

In the past century and a half since Pierre Paul Broca’s work with patient “Tan”, the study of language deficits in speakers with focal brain damage from stroke has been used to understand which brain regions are necessary for language. Recent applications of MRI analysis techniques allow better quantification of how focal brain damage impacts language deficits via lesion behavior mapping. The lesion behavior mapping approach provides critical evidence to clarify whether multiple regions identified in functional MRI studies are epiphenomenal or necessary for function. Here, for the first time we identify the brain regions required for producing words and organizing them during spontaneous connected speech before brain-behavior reorganization occurs in a large group of speakers with focal acute left hemisphere stroke.

Left hemisphere brain damage often impairs language production demonstrating that an extensive left frontal-temporal parietal network is necessary to produce multiple words in syntactically accurate structure (for reviews see Price, 2010; Fedorenko and Thompson-Schill, 2014; Hagoort and Indefrey, 2014). However, the processes for producing and organizing the content (i.e., words) into structure (lexical or syntactically accurate combinations) during connected speech localized within this left lateralized brain network are not well identified. This partly stems from a difficulty in eliciting spontaneous speech where deficits can be systematically quantified across individuals. As a result, almost all lesion behavior mapping analyses of connected speech in speakers with stroke have been limited to analyzing the content, i.e., the words elicited from picture descriptions or direct questions, or the fluency with which that content is produced (the number of words produced per minute or the number of words produced within utterances are often used as measures of the fluency of language production; Thompson *et al.*, 2013). In the first large lesion behavior mapping study of connected speech impairments, Borovsky *et al.* (2007) asked 50 subjects with diagnosed aphasia after left hemisphere stroke a series of autobiographical questions. Damage to a wide range of regions within frontal, parietal, and temporal lobes was associated with a reduced number of words produced overall and within utterances. Damage specific to posterior loci in the middle and superior temporal gyri as well as the angular gyri was associated with deficits in producing a reduced variety of words. However, the contribution of stroke severity as measured by lesion volume was not analyzed. Halai *et al.* (2017) examined similar aspects of connected speech during picture description using principal components analysis (PCA) in a smaller group of subjects (n = 31) with diagnosed aphasia after left hemisphere stroke (cf. Halai *et al.*, 2018). Here too, the ability to produce more words and more words quickly was associated with damage to frontal regions while reduced lexical diversity was associated with posterior regions including the middle and superior temporal gyri, and supramarginal gyrus. However, across these studies none of these regions survived correction for overall lesion volume. Using a multivariate lesion behavior mapping model and large left hemisphere chronic aphasia stroke sample (n = 90), Yourganov *et al.* (2016) found the speech fluency scores estimated as part of the Western Aphasia Battery (Kertesz, 1982) were predicted by damage to both left anterior (middle frontal gyrus and the inferior frontal gyrus – pars operculum) and posterior areas (supramarginal and posterior superior temporal gyri). However, here too lesion size predicted the fluency score and was highly correlated with the areas revealed. Significant findings controlling for the effect of lesion size were from a combined PCA and lesion behavior mapping analysis which included not only the connected speech measures but other production and cognitive measures. In these subsequent analyses, fluency related factors were primarily associated with the frontal lobe (Halai *et al.*, 2017; Lacey *et al.*, 2017; Halai *et al.*, 2018). Critically however, the fluency related PCA derived factors were comprised of additional variables with high loadings including naming, repetition, and reading among others which complicates the relationship between fluency deficits and brain damage localization. When controlling for lesion size, this relationship disappeared when performing the PCA within only connected speech measures (Halai *et al.*, 2017). In summary, although there appears to be an anterior (frontal) to posterior (temporal-parietal) division in terms of the number of words produced (fluency) vs. the ability to produce more diverse lexical content during spontaneous connected speech, the ability to produce content is best accounted for by stroke severity not the location of damage, i.e. the more brain damage incurred the fewer and less-diverse words produced, irrespective of where the damage occurred.

Relatively little is known about how individual differences in focal brain damage affect the ability to accurately produce syntactic structure independent from content. In the only lesion behavior mapping study of content and structure deficits during spontaneous connected speech of which we are aware, Mirman *et al.* (2019) examined story-telling narratives from 46 speakers with diagnosed aphasia after chronic left hemisphere stroke. After controlling for the contribution of lesion size, the proportion of words produced in sentences (utterances with both a subject noun and verb) was associated with damage to left middle and inferior frontal gyri, postcentral gyrus, and inferior parietal lobe. However, no lesion pattern emerged to predict syntactic deficits measured by the number of closed-class words (e.g., determiners, prepositions) produced out of all words in the narrative. No other aspects of spontaneous connected speech were examined. It remains unknown whether damage to focal regions within the left hemisphere language network critical for producing content dissociates from damage to regions responsible for generating syntactic structure during spontaneous connected speech.

### The current study

Here, we addressed the question of whether impairments producing the content and structure of spontaneously generated speech are related to different patterns of focal brain damage. Whether unique brain regions are required to produce content and structure during connected speech speaks to a central debate concerning the specificity of brain regions required for syntactic processing (e.g., Wilson and Saygın, 2004; Thothathiri *et al.*, 2012; Magnusdottir *et al.*, 2013; Blank *et al.*, 2016; Wilson *et al.*, 2016; Fedorenko *et al.*, 2018; Rogalsky *et al.*, 2018; cf. Dapretto and Bookheimer, 1999; Hagoort and Indefrey, 2014; Friederici *et al.*, 2017). To our knowledge, this is the first study of individual differences during the acute stage of stroke linking brain damage to deficits in the content and structure of spontaneous connected speech.

Our approach has significant methodological strengths. First, by examining lesion behavior relationships in a large group of speakers identified with radiological signs of left hemisphere acute stroke, we increased variability in behavioral performance, lesion size, and lesion location and avoided the confound of brain-behavior reorganization. Studies of speakers with chronic stroke often preselect subjects based on aphasia diagnosis (e.g. Schnur *et al.*, 2009; Yourganov *et al.*, 2016; Halai *et al.*, 2017; Mirman *et al.*, 2019) which limits behavioral and lesion location variability. Speakers with chronic aphasia typically have large lesions spanning crucial language areas (e.g. inferior frontal gyrus, posterior superior temporal gyrus and angular gyrus) where adjacent cortical regions are often damaged and damage is correlated with overall lesion size (Ochfeld *et al.*, 2010; Yourganov *et al.*, 2016; Shahid *et al.*, 2017). As a result, it is difficult to disentangle how lesion size and damage to specific regions contribute to function. Further, function may have shifted to other regions after reorganization occurred (Weiller *et al.*, 1995; Marsh and Hillis, 2006; Saur *et al.*, 2006; Thompson and den Ouden, 2008; Turkeltaub *et al.*, 2012; Nardo *et al.*, 2017; Hartwigsen and Saur, 2019). Here, we increased power to assess whether damage to a region significantly affected behavior in comparison to no damage to that region, by providing sufficient behavioral variability and heterogeneity for both between- and within-region lesion distributions (cf. Kimberg *et al.*, 2007; Sperber and Karnath, 2017; Lorca-Puls *et al.*, 2018; Pustina *et al.*, 2018; Sperber *et al.*, 2019). Second, we elicited speech using a spontaneous story-telling generation task which provides the advantages of eliciting grammatically diverse spontaneous speech (Thompson *et al.*, 2010b; den Ouden *et al.*, 2019) while providing excellent ecological validity with abilities to produce everyday speech (Olness and Ulatowska, 2017) and good test re-test reliability (Brookshire and Nicholas, 1994a, b; Roberts and Post, 2018). Third, we conducted detailed quantitative analyses to independently assess the content and structure of spontaneous speech using the rigorous quantitative production analysis approach (Saffran *et al.*, 1989; Rochon *et al.*, 2000; Gordon, 2006; Wilson *et al.*, 2010). Fourth, because single behavioral measures may consist of multiple cognitive components, we applied PCA to extract the underlying cognitive components across the quantitative production analysis measures (Rochon *et al.*, 2000; Butler *et al.*, 2014; Mirman *et al.*, 2015a; Lacey *et al.*, 2017). Lastly, we applied support vector regression based multivariate lesion behavior mapping which considers the pattern of all voxels as a single model to predict a behavioral outcome. In comparison to univariate approaches, multivariate lesion behavior mapping ameliorates limitations from differential lesion distribution across voxels, Type II error from applying statistical corrections across voxels and allows for interactions between different damaged areas to account for behavior (Mah *et al.*, 2014; Zhang *et al.*, 2014; Yourganov *et al.*, 2016; Pustina *et al.*, 2018; Sperber *et al.*, 2019).

## Materials and Methods

### Participants

Ninety acute stroke patients were consecutively recruited from the comprehensive stroke centers at the Memorial Hermann, Houston Methodist, and St. Luke hospitals in Houston, Texas as part of an ongoing project. Following previous studies (Kristinsson *et al.* in press; Karnath *et al.*, 2010; Corbetta *et al.*, 2015) we included participants diagnosed with an acute ischemic or parenchymal hemorrhagic left hemisphere stroke. Further, we included patients if they were native English speakers and had no history of other significant neurological diseases (e.g., dementia, schizophrenia) as assessed by the clinical neurological care team and subsequently documented in electronic medical records. Informed consent was approved by the Baylor College of Medicine Institutional Review Board.

For the current study, 25 patients were excluded because they were not able to complete the spontaneous connected speech task (n = 20) or they produced unintelligible speech (n = 5). Sixty-five of the remaining patients (35 males; 54 right-handed; 4 hemorrhagic) had sufficient language production to complete the spontaneous connected speech task with no severe apraxia of speech to preclude accurate connected speech scoring. Mean apraxia of speech score (subtest 5 of the Second Edition of the Apraxia Battery for Adults; (Dabul, 2000) was 1.2 (s.d. = 0.6; range = 1-3) where seven patients who did not complete this task were assessed using a picture description (picnic) and the story-telling task. Mean age and education were 61 (s.d. = 14; range = 20-85) and 14 (s.d. = 4; range = 6-33) years, respectively. To minimize effects of brain-behavior reorganization, we completed behavioral testing within an average of four days after stroke onset (s.d. = 3; range = 1-12 days).

The control group consisted of 13 non-brain damaged participants (3 male, 11 right-handed) matched in age and education with the patient group (|*t*|’s < 1.44, *p* values > 0.16). Mean age and education were 55 (s.d. = 14, range = 37-78) and 16 (s.d. = 3; range = 12-22) years respectively.

### Story-telling assessment

Patients viewed a picture book of “Cinderella” with printed text occluded at bedside for as long as they wished (Saffran *et al.*, 1989; Zingeser and Berndt, 1990; Rochon *et al.*, 2000; MacWhinney *et al.*, 2011; Rogalski *et al.*, 2011; Thompson *et al.*, 2012; Martin and Schnur, 2019). Then they closed the book and told the story in their own words. The experimenter encouraged participants to speak more if output was limited. Responses were recorded by a nearby digital device.

### Quantitative production analysis

Spontaneous connected speech narratives were transcribed and scored according to the procedures in the quantitative production analysis (QPA) training manual by two research assistants (Berdnt *et al.*, 2000). We chose the QPA because it uses an objective rating approach to comprehensively quantify production deficits by analyzing multiple measures at both structural and morphological levels of connected speech to identify differences across patients (Saffran *et al.*, 1989). As shown in a recent study, raters achieved high reliability for transcription and scoring (Martin and Schnur, 2019). We calculated thirteen QPA measures (see Table 1) and converted patients’ QPA scores to *z*-scores to reflect the degree of deficit using the distribution of control subject QPA performance.

**Table 1.**
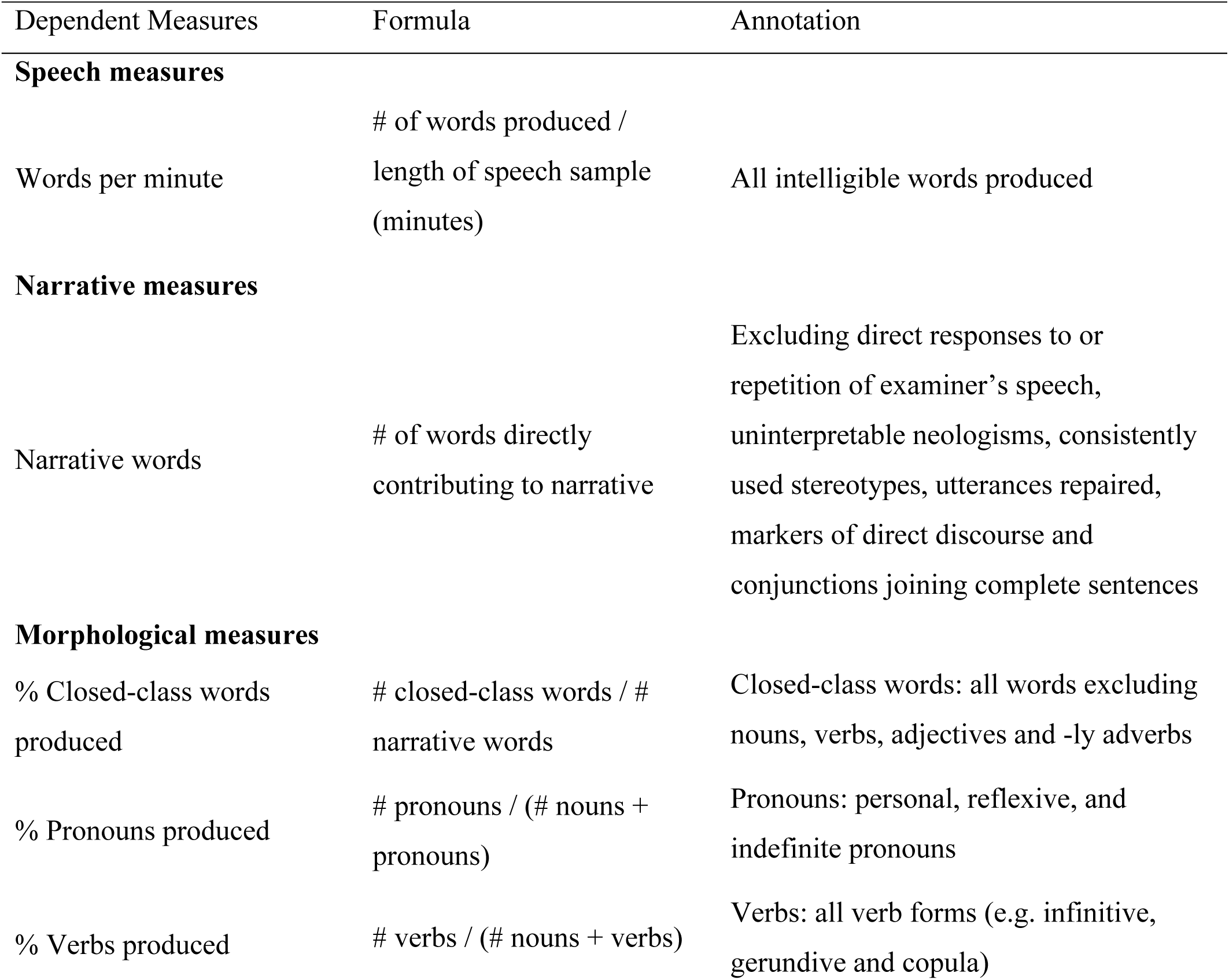
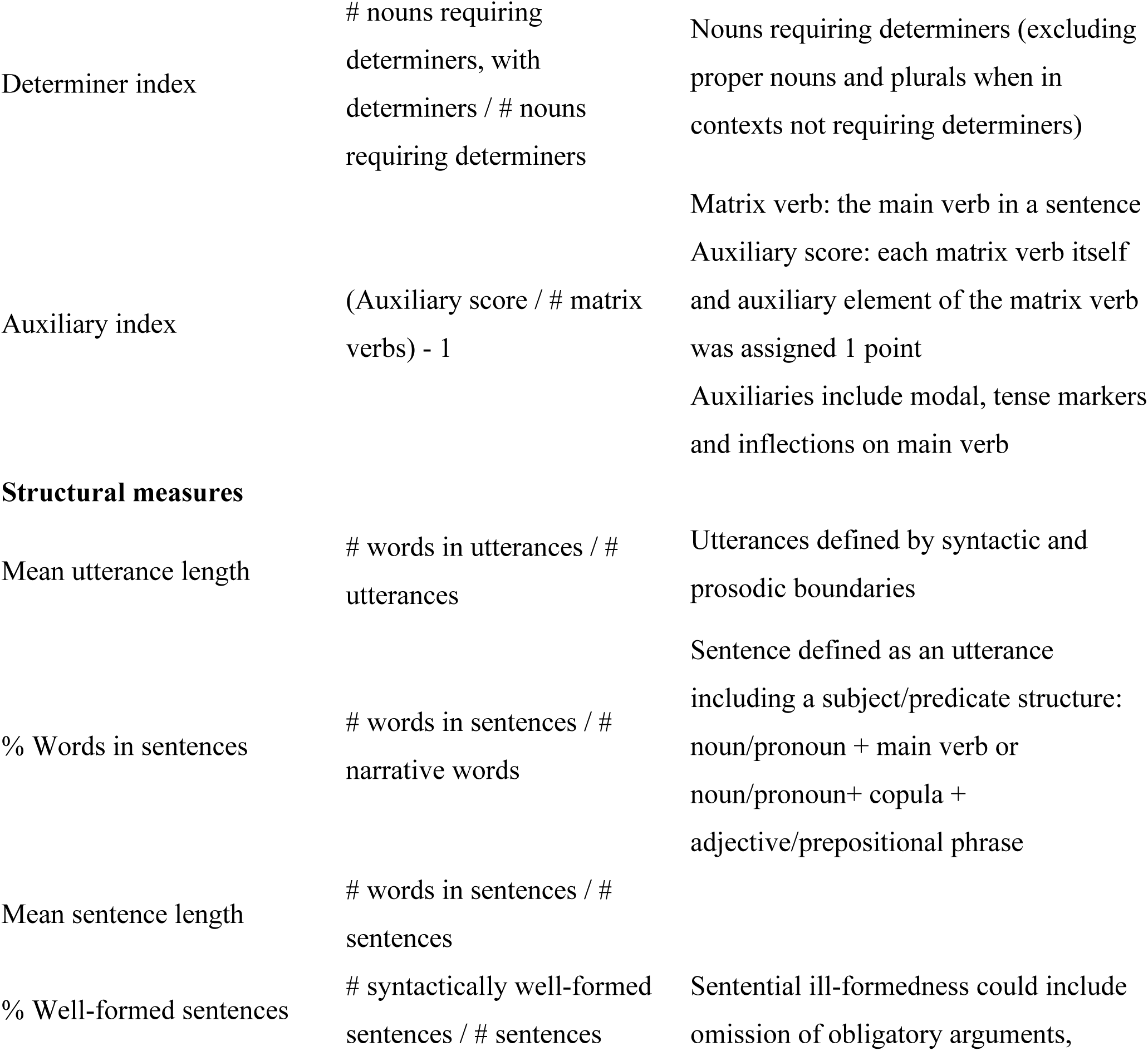
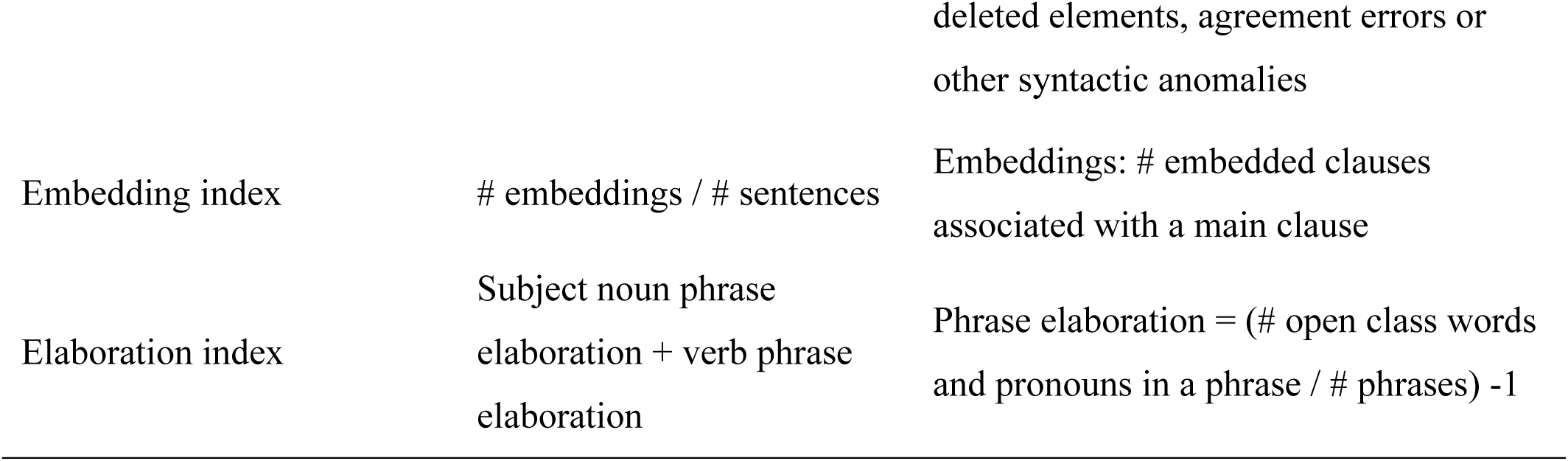
Summarized definitions of the quantitative production analysis measures following Saffran et al. (1989).

### Principle component analysis

To identify independent coherent subsets of the 13 quantitative production analysis measures of spontaneous connected speech, we used principal components analysis (PCA). Components with an eigenvalue exceeding one were extracted and rotated with the varimax method. To maximize power to detect significant effects for the PCA, we analyzed language performance for all subjects who fit selection criteria and were able to complete the connected speech task (n =65) independent of lesion demarcation constraints. We used varimax rotation because it only allows a small number of high variable loadings on each factor, yielding clearer interpretations (Kaiser, 1958). We calculated patient component scores for the lesion behavior mapping analysis.

### Imaging acquisition

We acquired diffusion weighted, apparent diffusion coefficient, and high-resolution structural scans (T1 and T2 FLAIR) along the axial direction with 4-5 mm slice thickness, and in cases where MRI was contraindicated, CT scans (n = 5) as part of the clinical protocols for admitted stroke cases. The voxel sizes of diffusion weighted and structural images were 1mm * 1mm * 4.5mm, and 0.5mm * 0.5mm * 4.5mm, respectively.

### Lesion tracing

To demarcate lesions, we first registered the diffusion weighted images with the high-resolution structural images (T1 or T2) using AFNI (https://afni.nimh.nih.gov/). Lesions were demarcated directly on the diffusion weighted images (by C.H), using ITK-snap (http://www.itksnap.org/pmwiki/pmwiki.php) with reference to apparent diffusion coefficient and T2 FLAIR images. Next, we normalized the individual structural images to the Colin-27 template in Montreal Neurological Institute space using ANTS registration (http://stnava.github.io/ANTs/; Avants *et al.*, 2008). We used the corresponding affine parameter and diffeomorphic maps to warp individual masks to the Montreal Neurological Institute space (Holmes *et al.*, 1998; Avants *et al.*, 2006). For patients with CT images but no MRI, lesions which were clearly visible were directly demarcated onto the Colin 27 template based on the CT image.

### Multivariate lesion behavior mapping analysis

To investigate lesion behavior relationships, we applied support vector regression (libsvm 3; https://www.csie.ntu.edu.tw/~cjlin/libsvm/) implemented with Matlab 2018b (https://www.mathworks.com/products/matlab.html). For lesion behavior mapping, eleven patients for whom lesions were difficult to identify were excluded [no clearly identifiable lesion (n = 7); subarachnoid hemorrhage (n = 3); and missing diffusion weighted imaging sequence (n = 1). Two patients were excluded because one or more component scores were identified as extreme outliers (beyond 3 s.d. from the patient average). Therefore, 52 patients were included in the lesion behavior mapping analysis (3 hemorrhagic; 2 with CT scans; 8 left handers). Only voxels with a lesion ratio exceeding 5% (at least 3 people) were included (Sperber and Karnath, 2017). Regarding a potential influence of lesion size, we residualized component scores by considering the effect of lesion size (cf. LESYMAP; https://github.com/dorianps/LESYMAP). Then, we normalized the residual scores into an interval of [0, 1] to keep the same scale with the binary lesion pattern [normalized score = (component score-minimal value) / (maximal value-minimal value)]. We selected a non-linear radial basis function kernel to build the model (Zhang *et al.*, 2014; Mirman *et al.*, 2015b) because it provided a better data fit in comparison to linear-models. Because gamma and c parameters could affect the model, to determine the optimal parameter pairs, we carried out a grid search on c (10^-2 – 10^9) and gamma (10^-9 – 10^3) for model selection (the same range as scikit-learn: https://scikit-learn.org/stable/index.html). Specifically, for each parameter pair, five-fold cross-validation was used to examine the model’s prediction accuracy. That is, data were split into five-folds, and each time, we used four-folds to train the model and predicted the remaining one. We averaged the mean squared errors between testing and predicted scores of the five folds to reflect the prediction accuracy. To assess model statistical inference, we further generated 1000 random models by permuting behavioral scores and compared the prediction accuracy from the original data to those from the random data. The *p*-value was the probability that the random models had a lower mean squared error than the original model. By comparing *p* values for all the parameter pairs, we determined the optimal pair of c and gamma and their significance (*p* < 0.05). We then explored the behavior-related lesion location. Following the approach of (Zhang *et al.*, 2014), the parametric values of the non-linear model were projected back to the original brain space, reflecting the statistical importance of the voxels to the model (i.e. the beta map). A similar permutation test for each voxel was further derived by shuffling behavioral scores 1000 times with the optimal model parameters to calculate random beta values associated with each voxel. As a result, the p-value of each voxel was generated by comparing random beta values with the original and thresholded at *p* < 0.05. To note, we did not apply multiple comparison correction because this is still a controversial issue (cf. Zhang *et al.*, 2014; Mirman *et al.*, 2015b). We reported significant voxel locations based on the Human Brain Connectome Atlas using a cluster threshold of > 100 voxels (Fan *et al.*, 2016).

### Data availability

The data are potentially available by request to T.T.S.

## Results

### Impairments of spontaneous connected speech

We calculated 13 aspects of spontaneous connected speech using quantitative production analysis (Saffran *et al.*, 1989; Rochon *et al.*, 2000; Gordon, 2006). Patients demonstrated wide variability and decreased performance in comparison to controls across most measures^1^ (see Figure 1). Patient performance was 1.5 s.d. away from controls on an average of 3.9 of the 13 measures (30%), with wide variability across individuals (s.d.: 20%; range 0-85%)^2^. As a group in comparison to controls, patients produced significantly fewer narrative words, required determiners, embeddings, well-formed sentences, proportionally more pronouns than nouns and an overall slower speech rate (|*t*| values > 3.20, *p* values < 0.002, Bonferroni corrected). At an individual level, 27 of the 65 patients (42%) scored more than 1.5 s.d. away from control performance on these measures (s.d.: 18%; range 14%-66%). Performance on other measures in comparison to controls was significantly impaired without correction for multiple comparisons. Specifically, patients produced proportionally more verbs compared to nouns, fewer elaborations, and shorter sentence and utterance lengths than controls (|*t*| values: 2.61-2.64, *p* values: 0.01-0.03). At an individual level, on average 19 of 65 patients (29%) scored more than 1.5 s.d. away from control performance on these measures (s.d.: 10%; range 19%-42%). There were no significant group differences in degree of auxiliary use, the proportion of closed-class words produced, and the number of words produced in sentences (|*t*| values < 0.60, *p* values > 0.55). However, at an individual level, on average 9 of 65 patients (14%) scored more than 1.5 s.d. away from control performance on these measures (s.d.: 6%; range 11%-22%). Therefore, as a whole, the patients were significantly impaired in comparison to controls on multiple aspects of spontaneous connected speech, with the sparing of few abilities.

**Figure 1.**
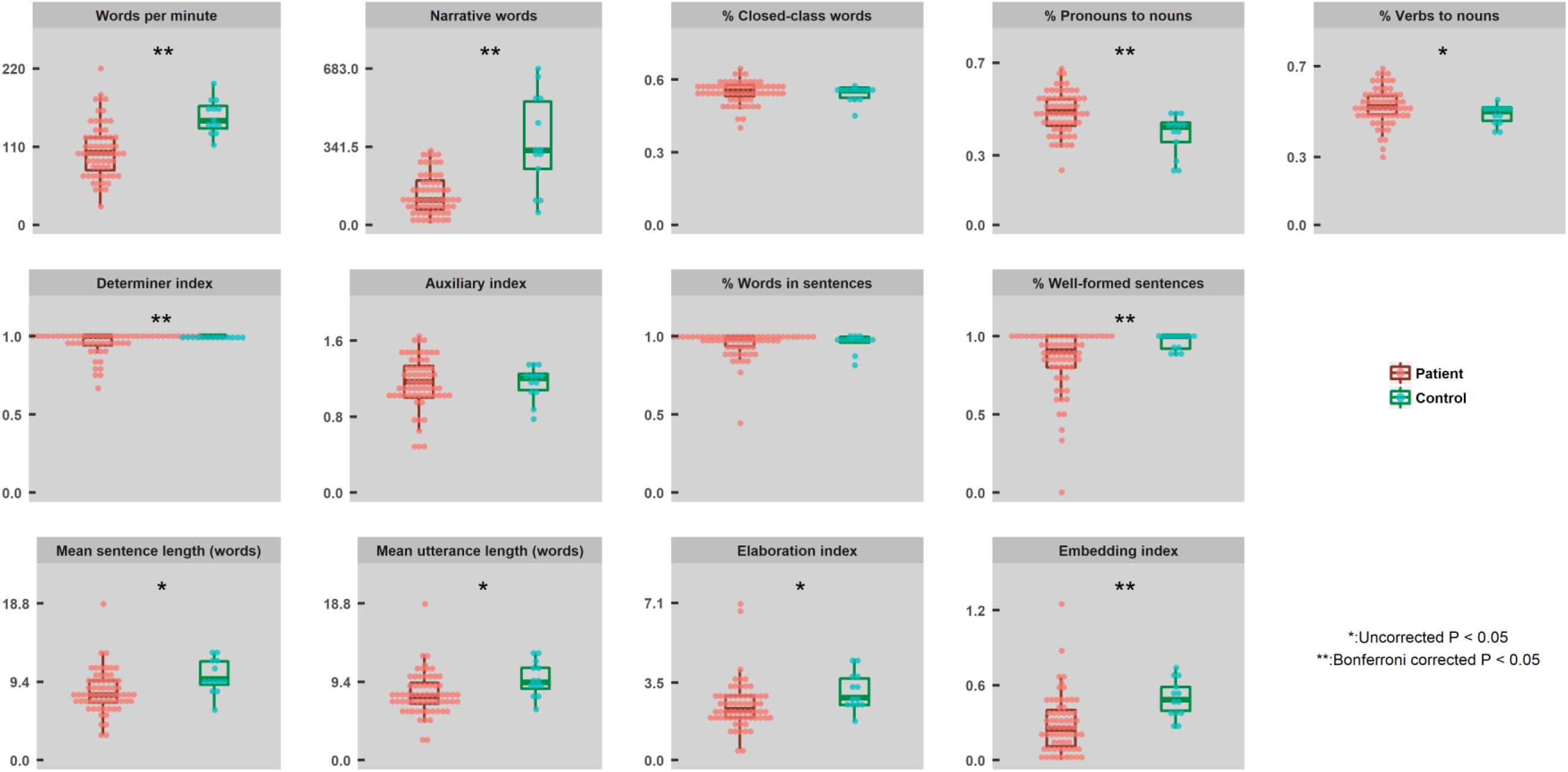
Individual performance (depicted by circles; patients-in red and controls-in green) and group distribution statistics (box and whisker plots depicting median, interquartile intervals, minimums, and maximums) for the quantitative production analysis measures of spontaneous connected speech. Significant group differences as determined by Welch’s unequal variances *t*-tests (Welch, 1947) indicated by **/*.

### Principal components of connected speech production

The quantitative production analysis measures of patients’ impairments during spontaneous connected speech production were appropriate for principal components analysis (PCA), as indicated by Kaiser-Meyer-Olkin (0.63) and Bartlett’s tests (*p* < 0.001; Pechenizkiy *et al.*, 2004). Using PCA, we extracted four connected speech components which accounted for 71% of the total variance, each component accounting for 30%, 16%, 15% and 10% of variance, respectively (see Figure 2). The first component had high loadings on the mean sentence length, mean utterance length, sentence elaboration index, embedding index and the number of narrative words produced (loadings > 0.6). We interpreted the first component as capturing the ability to produce words in increasingly complex combinations, which requires people to arrange words based on appropriate thematic and/or syntactic roles among other constraints. Hence, we call this component ‘structural complexity’. The second component had high loadings on the proportion of pronouns, proportion of verbs and proportion of closed-class words produced (loadings > 0.7). To note, this component is bidirectional. Negative values indicate difficulties in producing words with grammatical function, i.e. closed-class words, pronouns and verbs in comparison to primarily nouns. Positive values indicate the reverse. Thus, we interpret the second component as capturing ‘lexical selection’ abilities for different kinds of words (e.g., from nouns to closed-class words and verbs), depending on the individual subject’s component valence. The third component had high loadings on the proportion of well-formed sentences produced, proportion of words produced in sentences and the production of required determiners (loadings > 0.7). We refer to the third component as ‘syntactic accuracy’ as it captured the ability to generate accurate syntactic structure. The fourth component had high loading on a single variable, the number of words produced per minute (loading = 0.9). This component captured the fluency of overall spontaneous connected speech production. Hence, we referred to this component as ‘production fluency’. In summary, we characterized via PCA the patients’ multiple impairments of spontaneous connected speech production across four dimensions: structural complexity, lexical selection, syntactic accuracy and production fluency^3^

**Figure 2.**
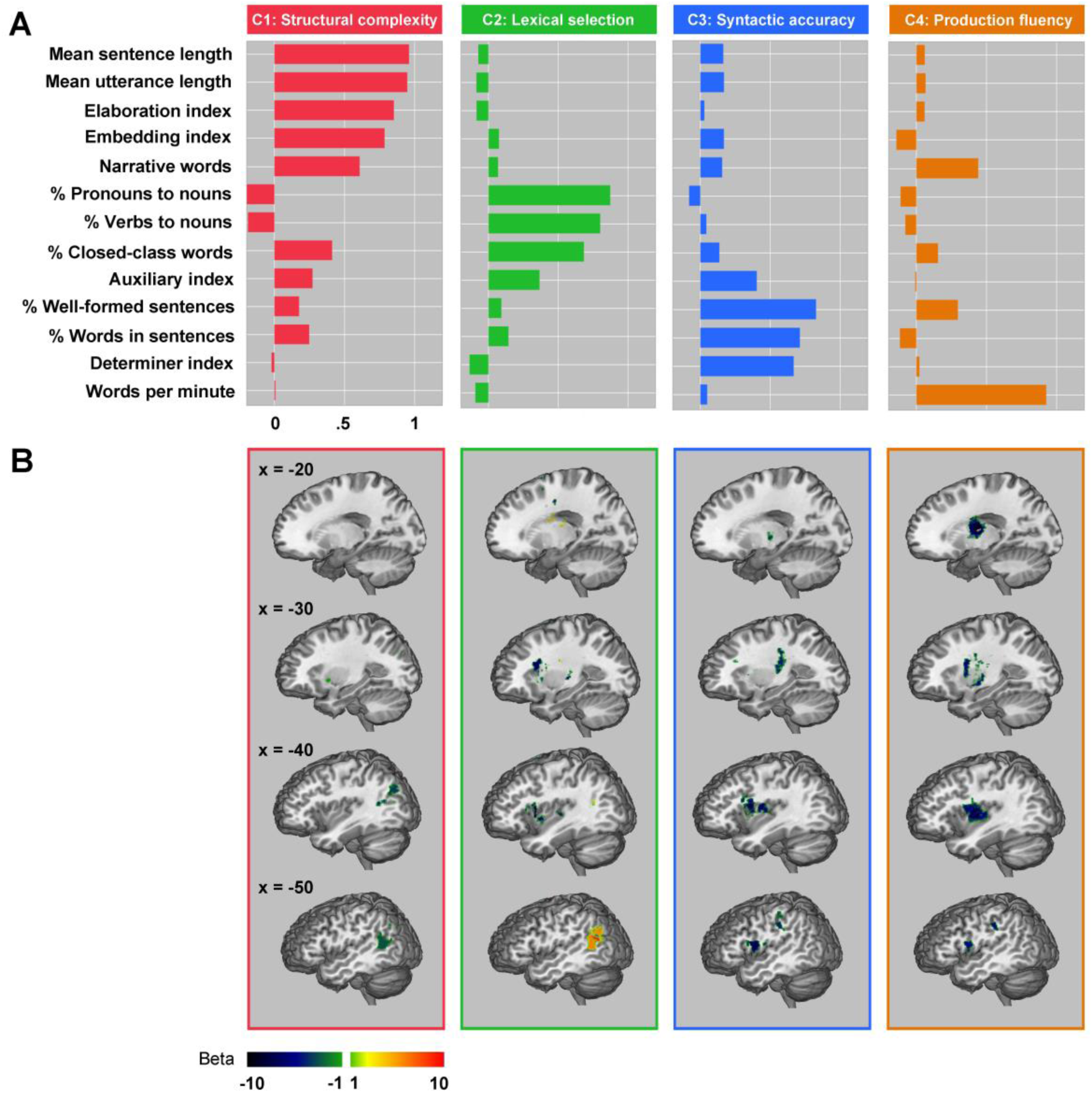
(A) Quantitative production analysis measure component loadings C1 – C4 and (B) significantly associated multivariate lesion behavior mapping beta maps.

### Lesion Distribution

We examined the distribution of voxels damaged in at least 5% (n = 3) of the patient cohort. Figure 3 displays the lesion distribution across subjects. The highest degree of lesion overlap was in the left basal ganglia (n = 11; 21% of the patient group; peak coordinates: −20, −14, 22). Other sufficiently lesioned voxels for lesion behavior mapping were distributed across the left middle and inferior frontal gyri, pre- and postcentral gyri, inferior and superior parietal lobes, posterior middle temporal gyrus, superior temporal sulcus, lateral occipital lobe, insula and thalamus.

**Figure 3.**
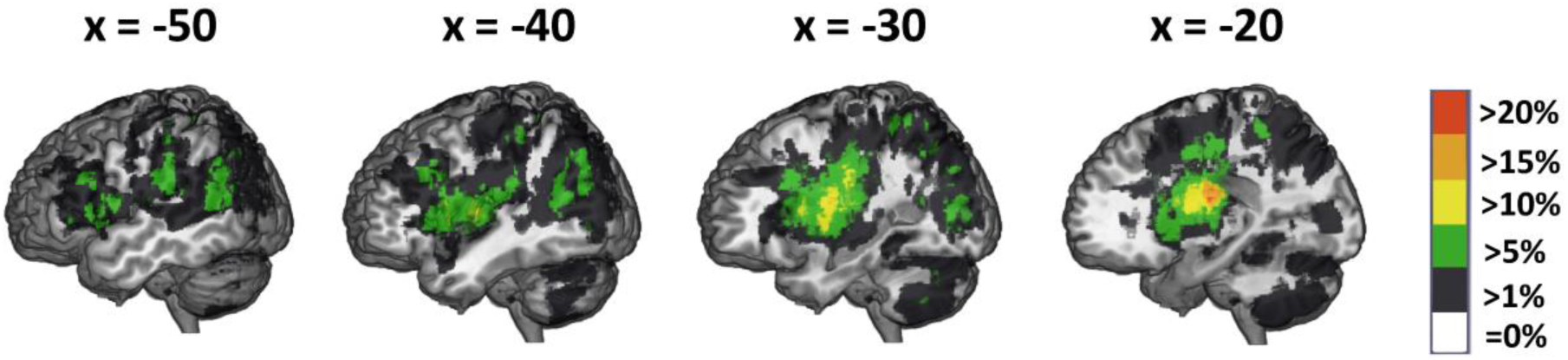
Lesion overlap across 52 patients. We conducted the lesion behavior mapping on voxels with damage in > 5% of the subject sample.

Acute lesions were on average smaller (our data: 12051 mm^3^) than typically seen in chronic stroke (reported average chronic lesion sizes vary between 21280 and 34176 mm^3^; cf. Corbetta *et al.*, 2015; Liew *et al.*, 2018), providing three distinct advantages for lesion behavior mapping. First, lesion volume was not significantly correlated with proportion damage to any brain connectome region (Bonferroni correction; average *r* = 0.23, range = −0.07-0.40; N.B. Regardless, we controlled for lesion volume in the lesion behavior mapping, see Section 2.4). This allowed us to examine the contribution of damage to function independent of overall stroke severity as measured by lesion volume. Second, regions farther apart from each other (Figure 4A, lighter green to white) were less likely to be damaged in the same individuals (e.g. inferior frontal gyrus and inferior parietal lobe), as illustrated by lower correlations in lighter red to white (Fig 4A) and thinner to no lines between regions (Fig 4B and 4C). The reduced cross-regional correlation in proportion damage within individuals in this cohort allowed us to tease apart the independent contribution of damage to behavior in frontal vs. temporal-parietal regions. Third, although most regions are relatively near each other (Fig 4A, darker green), proportion damage was correlated only between a few adjacent areas, and mostly within a brain connectome region (Fig 4A, darker red; Fig 4B and 4C, thicker lines between and within regions). As a result, we could discriminate between the contribution of damage to behavior for some adjacent areas (e.g. inferior frontal gyrus and insula).

**Figure 4.**
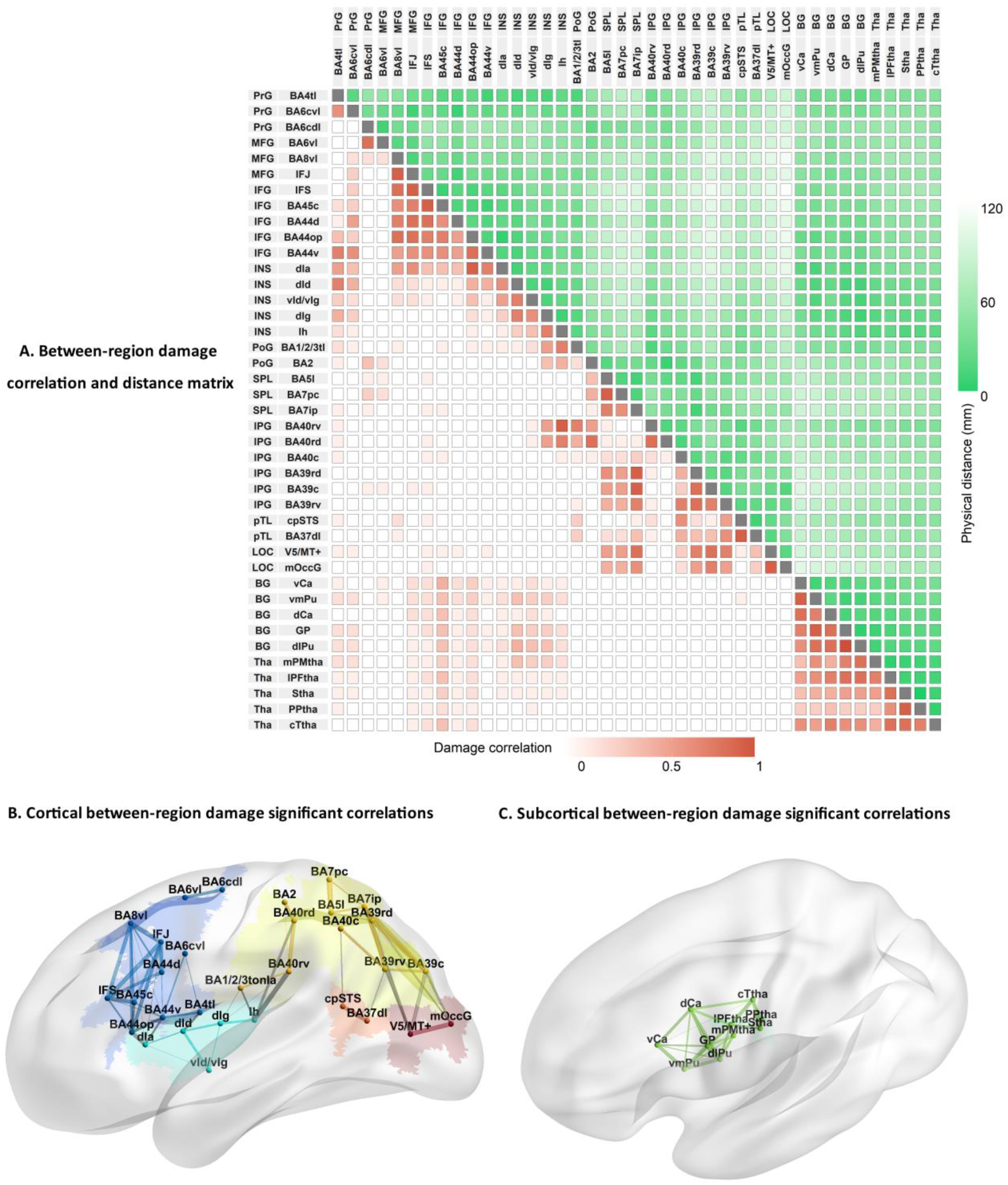
Proportion damage correlations between 41 brainconnectome regions (Fan *et al.*, 2016). A. Matrix of between region damage correlations and distances. Label colors depict different lobules: Blue: frontal lobe; Cyan: insula; yellow: parietal lobe; orange: temporal lobe; red: occipital lobe; green: thalamus. Increasing color intensity reflects either increasing correlations of proportion damage between regions (in red) or decreasing distance between regions (in green). Depictions of correlations between proportion damage across cortical (B) and subcortical regions (C). Lines’ thickness reflects the magnitude of correlation coefficients (where *r*’s > 0.53; *p* < 0.05 Bonfferroni correction). Correlations between lobes are shown in gray and within lobule as the same color as the lobule itself. MFG: middle frontal gyrus; IFG: inferior frontal gyrus; PrG: precentral gyrus; INS: insula; SPL: superior parietal lobe; IPL: inferior parietal lobe; PoG: postcentral gyrus; pTL: posterior temporal lobe; LOG: lateral occipital lobe; BG: basal ganglia; Tha: thalamus. IFJ: inferior frontal junction; IFS: inferior frontal sulcus; BA44op: opercular BA44; BA4tl: BA4 (tongue and larynx region); Ih: hypergranular insula; Ia: dorsal agranular insula; Ig: granular insula; Id: dysgranular insula; STS: superior temporal sulcus; BA7pc: postcentral BA7; BA7ip: intraparietal BA7; OccG: middle occipital gyrus; Ca: caudate; GP: globus pallidus; Pu: putamen: PMtha: pre-motor thalamus; Ptha: parietal thalamus; Ttha: temporal thalamus; PFtha: pre-frontal thalamus; c: caudal; r: rostral; d: dorsal; v: ventral; l: lateral; m: medial; s: superior; p: posterior.

### Neural substrates associated with impairments of spontaneous connected speech production

To explore lesion behavior relationships across different spontaneous connected speech production impairments, we conducted multivariate lesion behavior mapping using the support vector regression algorithm. For the lesion behavior mapping, we used the patients’ component scores and lesion masks. All the models were significant via cross-validation (*p* values < 0.03). Figure 2 and Table 2 illustrate the clusters with significant β values in each associated component model (voxel *p* < 0.05, cluster size > 100 voxels).

**Table 2.**
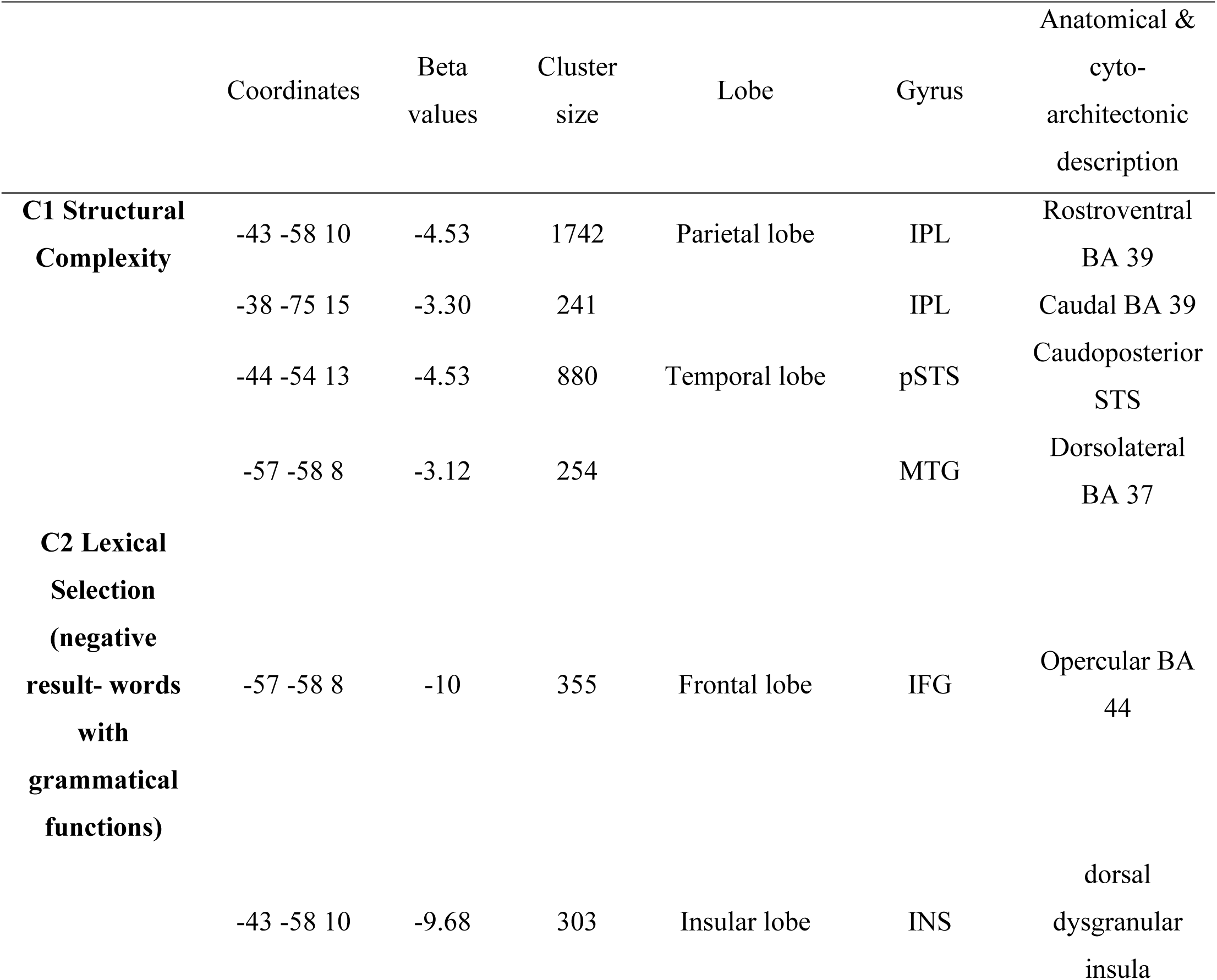
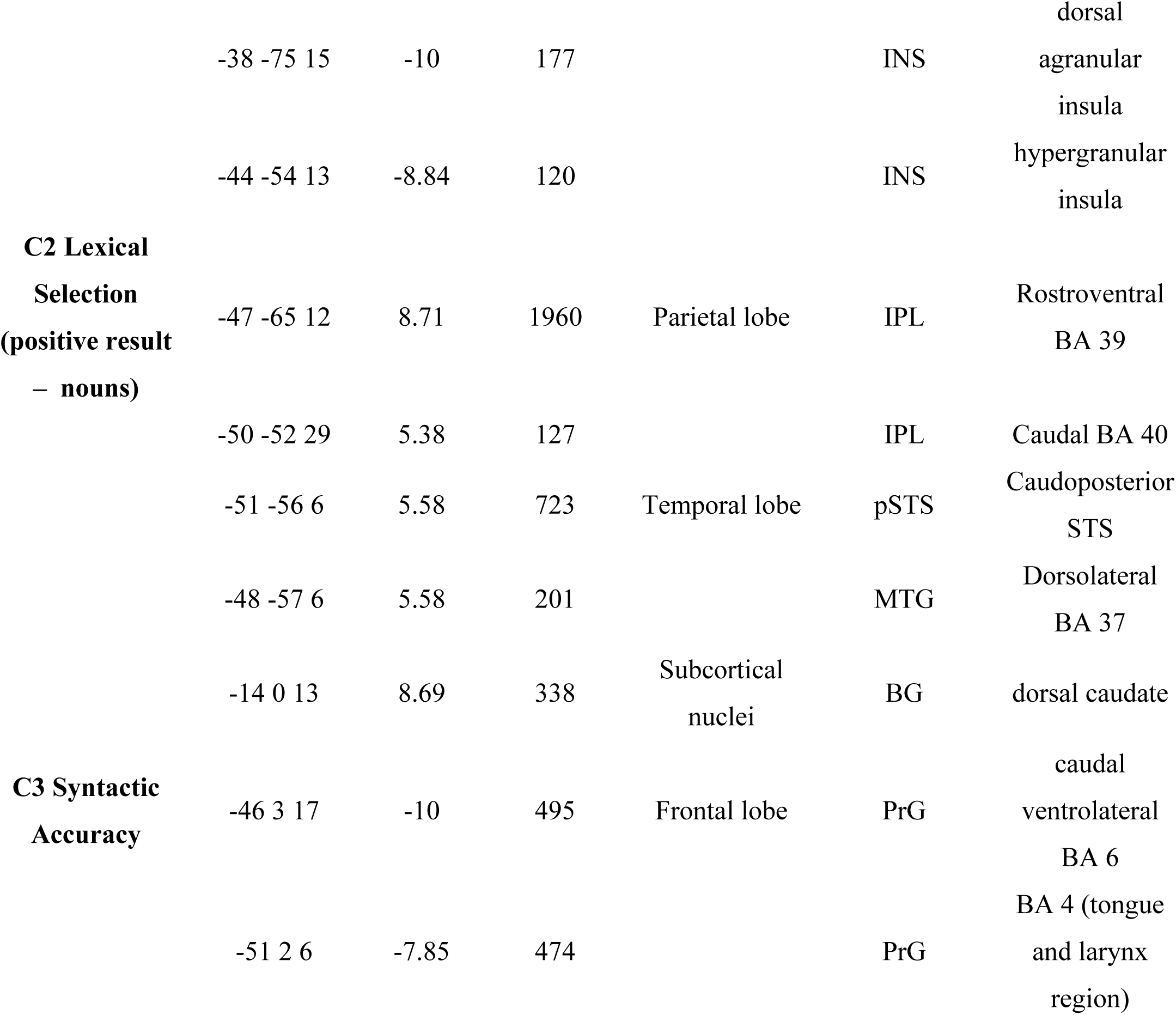
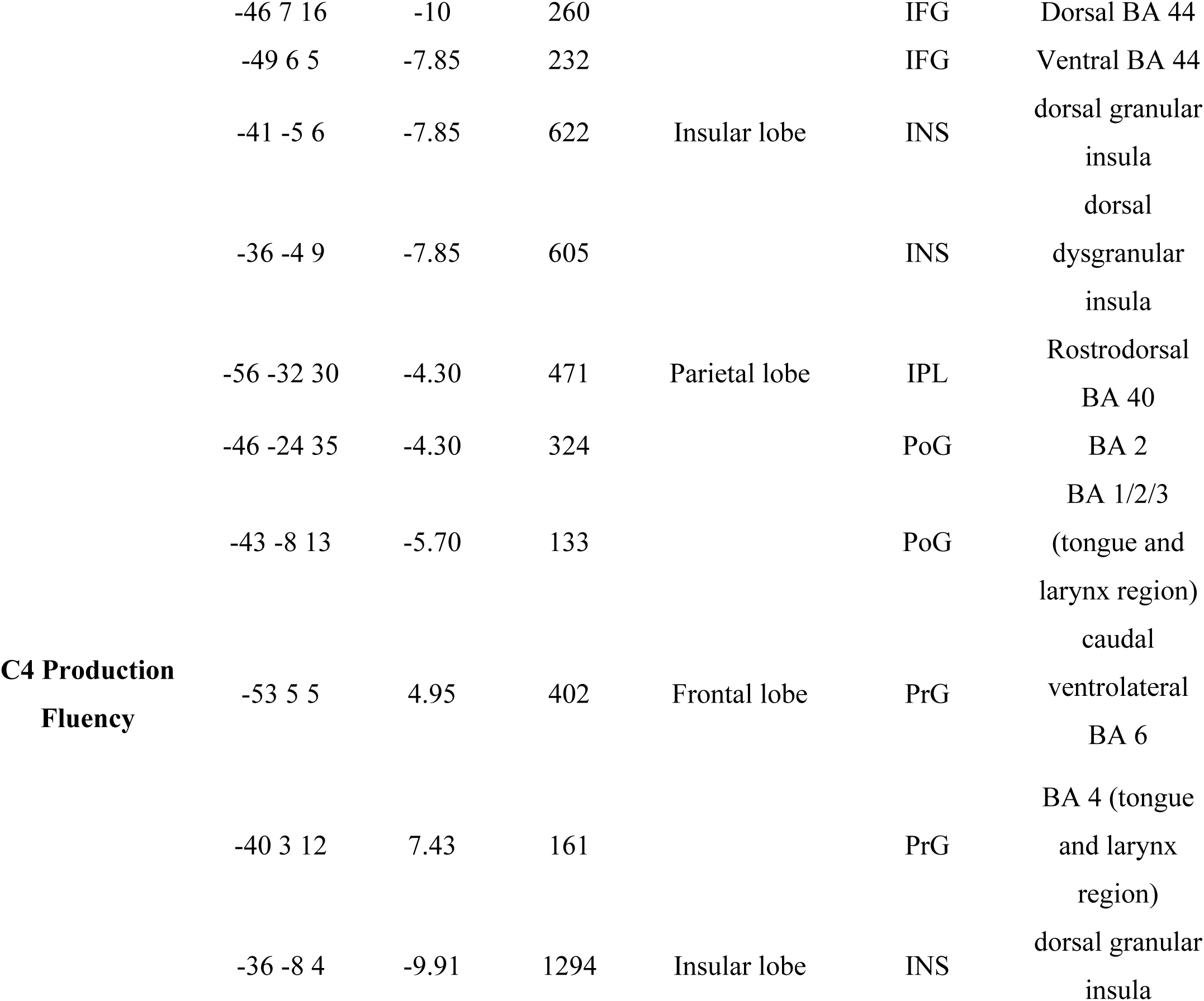
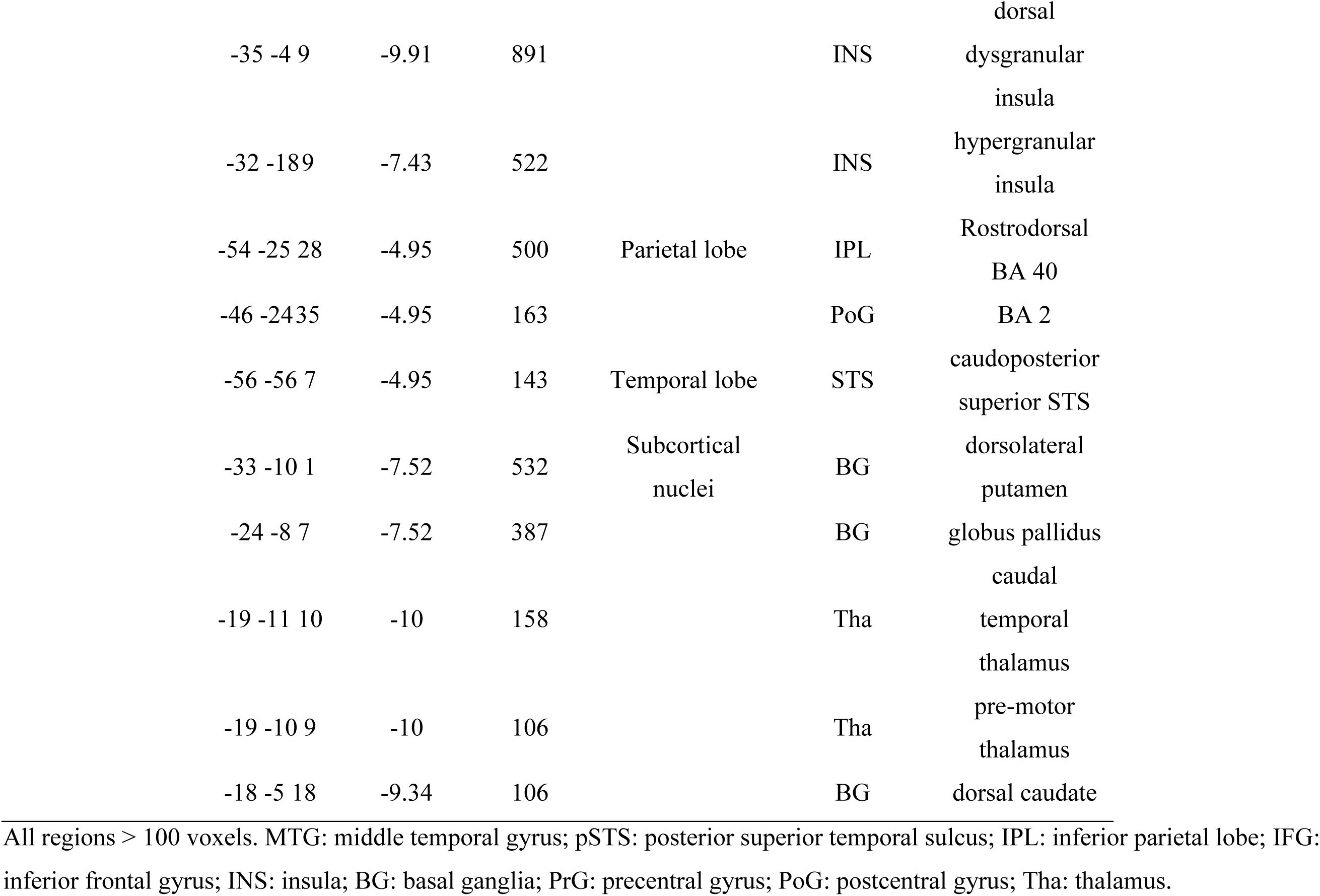
Regions significantly associated with the component score of speech production

The production of reduced structural complexity was associated with damage to areas within the left middle temporal gyrus, posterior superior temporal sulcus, and inferior parietal lobe. Impaired lexical selection (reduced production of nouns) was positively associated with damage to areas overlapping with those associated with reduced structural complexity as well as the basal ganglia. In contrast, impaired lexical selection specific to words with grammatical function (closed-class words, pronouns, and verbs) was negatively associated with damage to areas within the left inferior frontal gyrus and insula. Reduced syntactic accuracy was associated with damage to areas within the left inferior frontal gyrus, insula, precentral gyrus, postcentral gyrus and inferior parietal lobe. Impaired production fluency was associated with damage to multiple regions including the left precentral gyrus, postcentral gyrus, posterior superior temporal sulcus, inferior parietal lobe, insula, thalamus, and basal ganglia.

## Discussion

Here we asked whether producing words and organizing them into structure during spontaneous connected speech requires neuroanatomically and behaviorally distinct abilities. By quantifying brain damage and deficits in an unselected group of speakers during the acute phase of stroke we provide a detailed specification of the relationship between damage and deficits before brain-behavior reorganization occurs. Using PCA and multivariate lesion behavior mapping, our findings indicate that noun retrieval and the production of increasingly complex word combinations require exclusively left temporal-parietal regions while abilities to produce accurate syntactic structure and grammatical words (including verbs) are served primarily by left frontal regions. These results provide evidence from a rare and important patient population to clarify the debate as to whether morpho-syntactic processes are neuranatomically distinct from lexical processes required to produce spontaneous connected speech.

### Lexically-driven vs. syntactically-driven stages of production

A striking finding with important theoretical implications is the dissociation between damage to brain regions associated with the production of increasingly complex word combinations and damage associated with the production of syntactically accurate connected speech. Producing less complex structure (e.g., fewer words in utterances and phrases, fewer embedded clauses) was associated with damage to posterior temporal cortex [middle temporal gyrus (BA 37) and superior temporal sulcus] and the angular gyrus of the inferior parietal lobe (rostroventral BA 39)]. In contrast, syntactic accuracy deficits (e.g. omitting required noun determiners, verb arguments, or morphemes) were associated with damage to the left inferior frontal gyrus (dorsal and ventral BA 44; pars opercularis), the insula, precentral gyri (BAs 6 and 4), postcentral gyri (BAs 1-3), and a single locus in the supramarginal gyrus of the inferior parietal lobe (rostrodorsal BA 40).

We observed a similar rostral-caudal dissociation associated with deficits producing different types of words. Producing fewer nouns was associated with damage to overlapping regions associated with generating complex structure including the middle temporal gyrus (BA 37), posterior superior temporal sulcus, and the angular gyrus (BA 39). In contrast, producing fewer words with grammatical functions (e.g. determiners, prepositions, pronouns, and verbs) was associated with damage to regions near but not overlapping with those associated with syntactic accuracy deficits including BA 44 (pars opercularis) within the left inferior frontal gyrus and the insula.

We hypothesize that the dissociation between brain regions necessary for noun retrieval and their combination into increasingly complex structure vs. regions necessary for syntactic accuracy and grammatical word production reflect a neuroanatomical dissociation between different stages required to combine words into structure during connected speech. During initial stages of connected speech, a speaker generates a message she wishes to convey and activates the concepts associated with that message (Levelt, 1989; Levelt *et al.*, 1999). As Fromkin (1971) and Garrett (1975; 1980) originally formulated based on speech errors and others further developed (Dell, 1986; Levelt, 1989; Bock and Loebell, 1990; Bock and Levelt, 1994; Bock, 1995; Vigliocco and Hartsuiker, 2002; Matchin and Hickok, 2019), subsequent stages orchestrate word selection and the organization of words into larger multiword utterances. During functional encoding, a speaker must select the words associated with the concepts that form part of the intended message, potentially along with associated information like the word’s grammatical class (Levelt *et al.*, 1999; cf. Caramazza, 1997; Rapp and Goldrick, 2000) and for heads of phrases (nouns and verbs) the number of arguments (Vigliocco and Hartsuiker, 2002; Thompson *et al.*, 2010a; Kemmerer, 2015). To convey the event structure of the message, words are associated with each other via assignment of thematic roles (e.g. doer or receiver of an action) specified by the message and/or the lexical heads of phrase (cf. Vigliocco and Hartsuiker, 2002; Matchin *et al.*, 2019). Because thematic roles are not synonymous with syntactic functions (e.g. the agent of a message does not necessarily need to be the subject of the sentence, “The glass slipper was found by the prince”), elements are also assigned syntactic roles (e.g. subject, direct object; Thompson *et al.*, 2007; Thompson *et al.*, 2010a). We hypothesize that the middle temporal gyrus and posterior superior temporal sulcus are regions required for successful lexical selection and generation of lexically-driven argument structure (Saffran *et al.*, 1989; Bock and Levelt, 1994; Pallier *et al.*, 2011; Kemmerer, 2015). Interestingly, lexical selection and structural complexity associated regions were largely non-overlapping in the angular gyrus. An intriguing possibility based on comprehension (Wu *et al.*, 2007; Thothathiri *et al.*, 2012; cf. Thompson *et al.*, 2010a; Mack *et al.*, 2019) and production (den Ouden *et al.*, 2019) in stroke patients and fMRI evidence in unimpaired speakers (Pallier *et al.*, 2011; Matchin *et al.*, 2019; cf. Thompson *et al.*, 2007) is that the angular gyrus uniquely associated with structural complexity may be necessary for a conceptually (message driven), not lexically driven assignment of thematic roles (also see Binder and Desai, 2011). We speculate that these posterior regions are not involved in the assignment of syntactic roles because deficits in verb production (and syntactic accuracy) were not associated with their damage, consistent with proposals by Thompson and colleagues (2007, 2010; cf. Matchin et al., 2019). In sum, the overlap between temporal-parietal regions necessary for lexical selection and producing structural complexity suggests that these regions integrate content and structure corresponding to the functional encoding stage of connected speech production.

In contrast, we hypothesize that the regions associated with the production of accurate syntactic structure and grammatical words reflect a subsequent stage of grammatical encoding which positions utterance elements in order (Garrett, 1976; Bock and Levelt, 1994). This positional encoding stage involves the syntactic realization and linear ordering of lexical-semantic structures marked with bound- (e.g., inflectional markers indicating number, tense, and aspect, e.g., -ed in English) and free-standing closed-class morphemes (e.g., determiners, prepositions). Positional encoding may further involve the phonological specification of this linear order via the ordering of phonological segments (Garrett, 1975; Bock, 1987; Berndt *et al.*, 2002; Boeckx *et al.*, 2014) and the generation of prosodic structure (Vigliocco and Hartsuiker, 2002; cf. Ferreira, 1993). Although our analysis of connected speech did not quantify deficits at a phonological level of specificity, the postcentral and supramarginal gyri we identified are similar to damaged regions associated with phonological and phonetic errors during picture naming by speakers with chronic stroke (Mirman *et al.*, 2019). Damage to the left inferior frontal gyrus and insula was associated with deficits producing words with grammatical functions and nearby regions were also associated with reduced syntactic accuracy. We speculate that this association may reflect a verb-directed component of the positional assembly of lexical-semantic structures (cf. Hagoort, 2013; Takashima *et al.*, 2019). Together these patterns suggest that the regions associated with morpho-syntactic and grammatical word deficits are required for the syntactic realization of linearized lexical-semantic structure and potentially associated phonological processing during the positional stage of grammatical encoding.

### Fluency

The last significant finding concerns fluency, the ability to produce connected speech without effort. We identified an extensive frontal-temporal-parietal and subcortical network of brain regions associated with fluency. This is potentially unsurprising as fluency as reflected here by the number of words produced per minute likely depends on multiple processes during language production including message generation, conceptual activation, grammatical encoding, phonological, articulatory and motor processes (cf. Nozari and Faroqi-Shah, 2017). These results overlap but are not synonymous with regions associated with fluency via lesion behavior mapping in chronic stroke (Yourganov *et al.*, 2016; Halai *et al.*, 2017; Lacey *et al.*, 2017; Halai *et al.*, 2018; cf. Borovsky *et al.*, 2007) and brain atrophy in patients with primary progressive aphasia (Wilson *et al.*, 2010; Rogalski *et al.*, 2011). Differences in our approach to measuring fluency and the results extend previous work in several ways. First, we examined fluency quantitatively (cf. Yourganov *et al.*, 2016) and without contribution of other cognitive measures not reflective of connected speech (e.g. Lacey *et al.*, 2017; Halai *et al.*, 2018). Second, the relationship between fluency and associated brain regions was not due to overall lesion volume (e.g. Borovsky *et al.*, 2007; Yourganov *et al.*, 2016; Halai *et al.*, 2017, 2018) or diffuse damage (e.g. Wilson *et al.*, 2010; Rogalski *et al.*, 2011). Lastly, we examined spontaneous connected speech during story-telling as opposed to picture description (Wilson *et al.*, 2010; Yourganov *et al.*, 2016; Halai *et al.*, 2017; Lacey *et al.*, 2017; Halai *et al.*, 2018; cf. Borovsky *et al.*, 2007; Rogalski *et al.*, 2011), where speakers could not rely on visual input to guide message generation and concept identification. As a result, the fluency related brain network we identified is likely a more precise and ecologically relevant reflection of the multiple brain regions necessary for producing fluent connected speech.

### Implications

This study demonstrated neuroanatomical and functional divisions within grammatical encoding during connected speech which dovetail with existing neurobiological sentence production evidence. In chronic stroke speakers, lesion behavior mapping demonstrated that reduced numbers of words produced in sentences during story-telling was associated largely with damage to the left inferior frontal gyrus with smaller foci in the postcentral gyrus and inferior parietal lobe (Mirman *et al.*, 2019). When speakers undergoing awake craniotomies described pictures depicting actions, direct cortical stimulation of the pars opercularis and triangularis of the left inferior frontal gyrus resulted in 50% of patients producing grammatical errors during sentence production (Chang *et al.*, 2018). In speakers diagnosed with primary progressive aphasia, atrophy in primarily frontal regions was related with grammatical deficits during picture description (Wilson *et al.*, 2010). The role of these left inferior frontal regions is also consistent with regions implicated for patients clinically assessed with classic agrammatic aphasia profiles using standard aphasia batteries (Saffran *et al.*, 1989; den Ouden *et al.*, 2019; cf. Rochon *et al.*, 2000).

However, neural localization of syntactic processing during sentence comprehension is more equivocal. On the one hand, that the posterior portion of the left inferior frontal gyrus (BA 44) is associated with deficits to produce appropriate syntactic markers and closed-class elements is consistent with proposed divisions of labor for the left inferior frontal gyrus in sentence comprehension (for reviews of evidence see Hagoort and Indefrey, 2014; Friederici *et al.*, 2017; cf. Hickok and Poeppel, 2007). However, deficits related to syntactic processing in sentence comprehension are also associated with left temporal-parietal regions (Rogalski *et al.*, 2011; Thothathiri *et al.*, 2012; Magnusdottir *et al.*, 2013; Caplan *et al.*, 2016; Fridriksson *et al.*, 2018; cf. Tyler *et al.*, 2010; Rogalsky *et al.*, 2018) raising the question of the role of anterior regions in syntactic processing (e.g. Thothathiri *et al.*, 2012; Magnusdottir *et al.*, 2013; Blank *et al.*, 2016; Wilson *et al.*, 2016; Fedorenko *et al.*, 2018; Rogalsky *et al.*, 2018).

Matchin and Hickok (2019) propose that the inherently different processing requirements for language production and comprehension account for the differences in syntactic processing related neural substrates. In production as well as comprehension, the generation of lexical-semantic structure is required both to understand the relationship between words during comprehension, as well specify relationships between words to accurately reflect the message to be produced. However, only in production is morpho-syntactic (and phonological) processing necessary to order elements to accurately reflect the words and associated message to be conveyed. Matchin and Hickok advocate Ferreira and colleagues’ proposal (Christianson *et al.*, 2001; Ferreira *et al.*, 2002) that understanding who did what to whom during sentence comprehension can often proceed based on discourse-level and semantic information without decoding of morphosyntactic elements which may explain why anterior regions are not consistently engaged for syntactic processing during sentence comprehension. Our results significantly contribute to this theoretical debate by demonstrating that for spontaneous connected speech production, frontal and temporal-parietal regions are differentially necessary for lexically-driven (functional) vs. morpho-syntactic (positional) encoding.

### Limitations

Our study had limitations. First, because we examined connected speech deficits during the acute phase of stroke, individuals with minimal language output could not be included. This resulted in an underrepresentation of individuals with the most severe production impairments and potentially damage to additional regions within the left hemisphere language network (e.g. the anterior temporal lobe). This represents a trade-off between examining behavior before brain-behavior reorganization and avoiding lesion volume confounds vs. assessing more severe deficits. Second, compared with univariate approaches, multivariate lesion behavior mapping identifies multiple brain regions associated with function (Zhang *et al.*, 2014; Yourganov *et al.*, 2016). Within these brain region networks, not only the brain regions themselves but also their connections contribute to function. For example, impaired structural and functional connectivity of the arcuate fasciculus, which connects the inferior frontal gyrus and temporal parietal junction relates to multiple aspects of spontaneous speech production deficits in speakers with chronic stroke and primary progressive aphasia, such as informativeness, syntax or fluency (Marchina *et al.*, 2011; Wilson *et al.*, 2011; Yourganov *et al.*, 2018). However, the neuroimaging data available when we assessed behavior during the acute phase of stroke were not sufficient to perform structural or functional connectivity analyses. Relatedly, we likely underestimated the behavioral contribution of some regions because focal lesions induce decreases in neuronal activation in distant regions (i.e., diaschisis; Carrera and Tononi, 2014). To minimize these limitations in future work, we anticipate investigating the structural connections (using diffusion tensor imaging) and functional connections (using resting state fMRI) related with spontaneous speech deficits in individuals during the subacute phase of stroke, independent of their ability to produce speech acutely.

### Conclusions

This work provides the first evidence in acute stroke of a functional and anatomical dissociation between temporal-parietal and frontal focal brain regions for lexically and syntactically driven processes required for spontaneous connected speech. These results are consistent with predictions from models of syntactic processing based primarily on evidence from language comprehension (Friederici *et al.*, 2017; Hagoort, 2019; Matchin and Hickok, 2019; cf. Wilson *et al.*, 2016; Fedorenko *et al.*, 2018; Rogalsky *et al.*, 2018). By including subjects unbiased as to clinical diagnosis of aphasia and lesion location in the acute stage of left hemisphere stroke, lesion sizes were generally smaller and more varied than in chronic stroke studies. As a result, we were able to dissociate lesion behavior relationships between brain regions while minimizing the confound of overall lesion size and before brain behavior reorganization occurred. Future directions should explore the mechanisms by which the critical components of connected speech identified here recover after acute stroke and whether other cognitive capacities like working memory (cf. Martin and Schnur, 2019) play a role in recovery. The answers to these questions will be important to help with therapeutic interventions whereby improving other cognitive capacities may incidentally, and critically, improve language function thus contributing to improved daily quality of life after stroke.

## CRediT authorship contribution statement

Junhua Ding: Formal analysis, Methodology, Visualization, Writing – original draft.

Randi Martin: Formal analysis, Writing – review & editing, Funding acquisition.

Cris A. Hamilton: Formal analysis, Investigation.

Tatiana T. Schnur: Conceptualization, Methodology, Writing – original draft, Supervision, Funding acquisition.

## Acknowledgments

The authors wish to thank Chia-Ming Lei, Danielle Rossi, and Miranda Brenneman for data collection. We thank Eric Johns, Bowie Lin, Hao Yan, and Rachel Zahn for transcription and analysis of narrative speech samples. We thank Lynn Maher for her help in diagnosing apraxia of speech in our subject population. We thank the clinical neurological intensive care unit teams at the University of Texas Health Sciences Center and Memorial Hermann Hospital, The Houston Methodist Hospital, and the Baylor St. Luke’s Hospital for their assistance in patient recruitment and neurological assessment. We gratefully acknowledge and thank our research subjects and their caregivers for their willingness to participate in this research. This work was presented at the Academy of Aphasia in Macau, China (2019).

## Footnotes

1 Because years age and education between patient and control groups were not significantly different (Welch |*t*|’s < 1.24, *p*’s > 0.16), did not have unequal variance (Levine’s test F’s < 1.43, *p*’s > 0.24) and did not correlate with patient performance on any QPA variable (|*r*|’s < 0.19; *p*’s > 0.14; age was marginally correlated with words produced per minute, *r* = -.24, *p* = .053) we did not control for these variables when comparing patient vs. control connected speech performance.

2 The degree of speech deficit as measured by the quantitative production analysis variables was similar for the left- and right-handed patients. That is, left-handed patients (n=11) demonstrated connected speech deficits (at least 1.5 SD below controls) on an average of four quantitative production analysis variables (range 1 – 11) which was similar to right-handed patients’ performance (average = four variables, range = 0-10).

3 The principal components analysis (PCA) results differ in some respects in comparison to the PCA Rochon et al. (2000) conducted likely because the current PCA involved different variables and more than twice as many subjects who were unselected for aphasia diagnosis during the acute (vs. chronic) stage of stroke.

## References

Avants BB, Epstein CL, Grossman M, Gee JC. Symmetric diffeomorphic image registration with cross-correlation: Evaluating automated labeling of elderly and neurodegenerative brain. Medical Image Analysis 2008; 12(1): 26–41.

Avants BB, Schoenemann PT, Gee JC. Lagrangian frame diffeomorphic image registration: Morphometric comparison of human and chimpanzee cortex. Medical Image Analysis 2006; 10(3): 397–412.

Berdnt RS, Wayland S, Rochon E, Safffran E, Schwartz M. Quantitative production analysis : a training manual for the analysis of aphasic sentence production. Hove, East Sussex, UK: Psychology Press; 2000.

Berndt RS, Haendiges AN, Burton MW, Mitchum CC. Grammatical class and imageability in aphasic word production: their effects are independent. Journal of Neurolinguistics 2002; 15(3): 353–71.

Binder JR, Desai RH. The neurobiology of semantic memory. Trends in Cognitive Sciences 2011; 15(11): 527–36.

Blank I, Balewski Z, Mahowald K, Fedorenko E. Syntactic processing is distributed across the language system. NeuroImage 2016; 127: 307–23.

Bock K. An effect of the accessibility of word forms on sentence structures. Journal of Memory and Language 1987; 26(2): 119–37.

Bock K. Producing Agreement. Current Directions in Psychological Science 1995; 4(2): 56–61.

Bock K, Levelt W. Language production: Grammatical encoding. Handbook of psycholinguistics. San Diego, CA, US: Academic Press; 1994. p. 945–84.

Bock K, Loebell H. Framing sentences. Cognition 1990; 35(1): 1–39.

Boeckx C, Martinez-Alvarez A, Leivada E. The functional neuroanatomy of serial order in language. Journal of Neurolinguistics 2014; 32: 1–15.

Borovsky A, Saygin AP, Bates E, Dronkers N. Lesion correlates of conversational speech production deficits. Neuropsychologia 2007; 45(11): 2525–33.

Brookshire RH, Nicholas LE. Speech Sample Size and Test-Retest Stability of Connected Speech Measures for Adults With Aphasia. Journal of Speech, Language, and Hearing Research 1994a; 37(2): 399–407.

Brookshire RH, Nicholas LE. Test-Retest Stability of Measures of Connected Speech in Aphasia. Clinical Aphasiology 1994b; 22: 119–33.

Butler RA, Lambon Ralph MA, Woollams AM. Capturing multidimensionality in stroke aphasia: mapping principal behavioural components to neural structures. Brain 2014; 137(12): 3248–66.

Caplan D, Michaud J, Hufford R, Makris N. Deficit-lesion correlations in syntactic comprehension in aphasia. Brain and Language 2016; 152: 14–27.

Caramazza A. How Many Levels of Processing Are There in Lexical Access? Cognitive Neuropsychology 1997; 14(1): 177–208.

Carrera E, Tononi G. Diaschisis: past, present, future. Brain 2014; 137(9): 2408–22.

Chang EF, Kurteff G, Wilson SM. Selective Interference with Syntactic Encoding during Sentence Production by Direct Electrocortical Stimulation of the Inferior Frontal Gyrus. Journal of Cognitive Neuroscience 2018; 30(3): 411–20.

Christianson K, Hollingworth A, Halliwell JF, Ferreira F. Thematic Roles Assigned along the Garden Path Linger. Cognitive Psychology 2001; 42(4): 368–407.

Corbetta M, Ramsey L, Callejas A, Baldassarre A, Hacker Carl D, Siegel Joshua S, et al. Common Behavioral Clusters and Subcortical Anatomy in Stroke. Neuron 2015; 85(5): 927–41.

Dabul B. Apraxia Battery for Adults (2nd ed..). Austin, TX: PRO-ED Inc; 2000.

Dapretto M, Bookheimer SY. Form and Content: Dissociating Syntax and Semantics in Sentence Comprehension. Neuron 1999; 24(2): 427–32.

Dell GS. A spreading-activation theory of retrieval in sentence production. Psychological Review 1986; 93(3): 283–321.

den Ouden D-B, Malyutina S, Basilakos A, Bonilha L, Gleichgerrcht E, Yourganov G, et al. Cortical and structural-connectivity damage correlated with impaired syntactic processing in aphasia. Human Brain Mapping 2019; 40(7): 2153–73.

Fan L, Li H, Zhuo J, Zhang Y, Wang J, Chen L, et al. The Human Brainnetome Atlas: A New Brain Atlas Based on Connectional Architecture. Cerebral Cortex 2016; 26(8): 3508–26.

Fedorenko E, Mineroff Z, Siegelman M, Blank I. Word meanings and sentence structure recruit the same set of fronto-temporal regions during comprehension. bioRxiv 2018: 477851.

Fedorenko E, Thompson-Schill SL. Reworking the language network. Trends in Cognitive Sciences 2014; 18(3): 120–6.

Ferreira F. Creation of prosody during sentence production. Psychological Review 1993; 100(2): 233–53.

Ferreira F, Bailey KGD, Ferraro V. Good-Enough Representations in Language Comprehension. Current Directions in Psychological Science 2002; 11(1): 11–5.

Fridriksson J, den Ouden D-B, Hillis AE, Hickok G, Rorden C, Basilakos A, et al. Anatomy of aphasia revisited. Brain 2018; 141(3): 848–62.

Friederici AD, Chomsky N, Berwick RC, Moro A, Bolhuis JJ. Language, mind and brain. Nature Human Behaviour 2017; 1(10): 713–22.

Fromkin VA. The Non-Anomalous Nature of Anomalous Utterances. Language 1971; 47(1): 27–52.

Garrett M. Levels of processing in sentence production. Language production Vol 1: Speech and talk. London: Academic Press; 1980. p. 177–220.

Garrett MF. The Analysis of Sentence Production. In: Bower GH, editor. Psychology of Learning and Motivation. New York: Academic Press; 1975. p. 133–77.

Garrett MF. Syntactic processes in sentence production. New approaches to language mechanisms 1976; 30: 231–56.

Gordon JK. A quantitative production analysis of picture description. Aphasiology 2006; 20(2-4): 188–204.

Hagoort P. MUC (Memory, Unification, Control) and beyond. Frontiers in Psychology 2013; 4(416).

Hagoort P. The neurobiology of language beyond single-word processing. Science 2019; 366(6461): 55–8.

Hagoort P, Indefrey P. The Neurobiology of Language Beyond Single Words. Annual Review of Neuroscience 2014; 37(1): 347–62.

Halai AD, Woollams AM, Lambon Ralph MA. Using principal component analysis to capture individual differences within a unified neuropsychological model of chronic post-stroke aphasia: Revealing the unique neural correlates of speech fluency, phonology and semantics. Cortex 2017; 86: 275–89.

Halai AD, Woollams AM, Lambon Ralph MA. Triangulation of language-cognitive impairments, naming errors and their neural bases post-stroke. NeuroImage: Clinical 2018; 17: 465–73.

Hartwigsen G, Saur D. Neuroimaging of stroke recovery from aphasia – Insights into plasticity of the human language network. NeuroImage 2019; 190: 14–31.

Hickok G, Poeppel D. The cortical organization of speech processing. Nature Reviews Neuroscience 2007; 8(5): 393–402.

Holmes CJ, Hoge R, Collins L, Woods R, Toga AW, Evans AC. Enhancement of MR Images Using Registration for Signal Averaging. Journal of Computer Assisted Tomography 1998; 22(2): 324–33.

Kaiser HF. The varimax criterion for analytic rotation in factor analysis. Psychometrika 1958; 23(3): 187–200.

Karnath H-O, Rennig J, Johannsen L, Rorden C. The anatomy underlying acute versus chronic spatial neglect: a longitudinal study. Brain 2010; 134(3): 903–12.

Kemmerer D. Are the motor features of verb meanings represented in the precentral motor cortices? Yes, but within the context of a flexible, multilevel architecture for conceptual knowledge. Psychonomic Bulletin & Review 2015; 22(4): 1068–75.

Kertesz A. Western aphasia battery test manual. San Antonio, TX: Psychological Corp; 1982.

Kimberg DY, Coslett HB, Schwartz MF. Power in Voxel-based Lesion-Symptom Mapping. Journal of Cognitive Neuroscience 2007; 19(7): 1067–80.

Kristinsson S, Thors H, Yourganov G, Magnusdottir S, Hjaltason H, Stark BC, et al. Brain Damage Associated with Impaired Sentence Processing in Acute Aphasia. Journal of Cognitive Neuroscience In press; 0(0): 1–16.

Lacey EH, Skipper-Kallal LM, Xing S, Fama ME, Turkeltaub PE. Mapping Common Aphasia Assessments to Underlying Cognitive Processes and Their Neural Substrates. Neurorehabilitation and Neural Repair 2017; 31(5): 442–50.

Levelt WJ. Speaking: From intention to articulation. Cambridge, MA: The MIT Press; 1989.

Levelt WJM, Roelofs A, Meyer AS. A theory of lexical access in speech production. Behavioral and Brain Sciences 1999; 22(1): 1–38.

Liew S-L, Anglin JM, Banks NW, Sondag M, Ito KL, Kim H, et al. A large, open source dataset of stroke anatomical brain images and manual lesion segmentations. Scientific Data 2018; 5: 180011.

Lorca-Puls DL, Gajardo-Vidal A, White J, Seghier ML, Leff AP, Green DW, et al. The impact of sample size on the reproducibility of voxel-based lesion-deficit mappings. Neuropsychologia 2018; 115: 101–11.

Mack JE, Mesulam MM, Rogalski EJ, Thompson CK. Verb-argument integration in primary progressive aphasia: Real-time argument access and selection. Neuropsychologia 2019; 134: 107192.

MacWhinney B, Fromm D, Forbes M, Holland A. AphasiaBank: Methods for studying discourse. Aphasiology 2011; 25(11): 1286–307.

Magnusdottir S, Fillmore P, den Ouden DB, Hjaltason H, Rorden C, Kjartansson O, et al. Damage to left anterior temporal cortex predicts impairment of complex syntactic processing: A lesion-symptom mapping study. Human Brain Mapping 2013; 34(10): 2715–23.

Mah Y-H, Husain M, Rees G, Nachev P. Human brain lesion-deficit inference remapped. Brain 2014; 137(9): 2522–31.

Marchina S, Zhu Lin L, Norton A, Zipse L, Wan Catherine Y, Schlaug G. Impairment of Speech Production Predicted by Lesion Load of the Left Arcuate Fasciculus. Stroke 2011; 42(8): 2251–6.

Marsh EB, Hillis AE. Recovery from aphasia following brain injury: the role of reorganization. In: Møller AR, editor. Progress in Brain Research: Elsevier; 2006. p. 143–56.

Martin RC, Schnur TT. Independent contributions of semantic and phonological working memory to spontaneous speech in acute stroke. Cortex 2019; 112: 58–68.

Matchin W, Hickok G. The cortical organization of syntax. PsyArXiv 2019.

Matchin W, Liao C-H, Gaston P, Lau E. Same words, different structures: An fMRI investigation of argument relations and the angular gyrus. Neuropsychologia 2019; 125: 116–28.

Mirman D, Chen Q, Zhang Y, Wang Z, Faseyitan OK, Coslett HB, et al. Neural organization of spoken language revealed by lesion–symptom mapping. Nature Communications 2015a; 6: 6762.

Mirman D, Kraft AE, Harvey DY, Brecher AR, Schwartz MF. Mapping articulatory and grammatical subcomponents of fluency deficits in post-stroke aphasia. Cognitive, Affective, & Behavioral Neuroscience 2019.

Mirman D, Zhang Y, Wang Z, Coslett HB, Schwartz MF. The ins and outs of meaning: Behavioral and neuroanatomical dissociation of semantically-driven word retrieval and multimodal semantic recognition in aphasia. Neuropsychologia 2015b; 76: 208–19.

Nardo D, Holland R, Leff AP, Price CJ, Crinion JT. Less is more: neural mechanisms underlying anomia treatment in chronic aphasic patients. Brain 2017; 140(11): 3039–54.

Nozari N, Faroqi-Shah Y. Investigating the origin of nonfluency in aphasia: A path modeling approach to neuropsychology. Cortex 2017; 95: 119–35.

Ochfeld E, Newhart M, Molitoris J, Leigh R, Cloutman L, Davis C, et al. Ischemia in Broca Area Is Associated With Broca Aphasia More Reliably in Acute Than in Chronic Stroke. Stroke 2010; 41(2): 325–30.

Olness GS, Ulatowska HK. Aphasias. In: Cummings L, editor. Research in Clinical Pragmatics. Cham: Springer International Publishing; 2017. p. 211–42.

Pallier C, Devauchelle A-D, Dehaene S. Cortical representation of the constituent structure of sentences. Proceedings of the National Academy of Sciences 2011; 108(6): 2522–7.

Pechenizkiy M, Tsymbal A, Puuronen S. PCA-based feature transformation for classification: issues in medical diagnostics. Proceedings 17th IEEE Symposium on Computer-Based Medical Systems; 2004 25-25 June 2004; 2004. p. 535–40.

Price CJ. The anatomy of language: a review of 100 fMRI studies published in 2009. Annals of the New York Academy of Sciences 2010; 1191(1): 62–88.

Pustina D, Avants B, Faseyitan OK, Medaglia JD, Coslett HB. Improved accuracy of lesion to symptom mapping with multivariate sparse canonical correlations. Neuropsychologia 2018; 115: 154–66.

Rapp B, Goldrick M. Discreteness and interactivity in spoken word production. Psychological Review 2000; 107(3): 460–99.

Roberts A, Post D. Information Content and Efficiency in the Spoken Discourse of Individuals With Parkinson’s Disease. Journal of Speech, Language, and Hearing Research 2018; 61(9): 2259–74.

Rochon E, Saffran EM, Berndt RS, Schwartz MF. Quantitative Analysis of Aphasic Sentence Production: Further Development and New Data. Brain and Language 2000; 72(3): 193–218.

Rogalski E, Cobia D, Harrison TM, Wieneke C, Thompson CK, Weintraub S, et al. Anatomy of Language Impairments in Primary Progressive Aphasia. The Journal of Neuroscience 2011; 31(9): 3344–50.

Rogalsky C, LaCroix AN, Chen K-H, Anderson SW, Damasio H, Love T, et al. The Neurobiology of Agrammatic Sentence Comprehension: A Lesion Study. Journal of Cognitive Neuroscience 2018; 30(2): 234–55.

Saffran EM, Berndt RS, Schwartz MF. The quantitative analysis of agrammatic production: Procedure and data. Brain and Language 1989; 37(3): 440–79.

Saur D, Lange R, Baumgaertner A, Schraknepper V, Willmes K, Rijntjes M, et al. Dynamics of language reorganization after stroke. Brain 2006; 129(6): 1371–84.

Schnur TT, Schwartz MF, Kimberg DY, Hirshorn E, Coslett HB, Thompson-Schill SL. Localizing interference during naming: Convergent neuroimaging and neuropsychological evidence for the function of Broca’s area. Proceedings of the National Academy of Sciences 2009; 106(1): 322–7.

Shahid H, Sebastian R, Schnur TT, Hanayik T, Wright A, Tippett DC, et al. Important considerations in lesion-symptom mapping: Illustrations from studies of word comprehension. Human Brain Mapping 2017; 38(6): 2990–3000.

Sperber C, Karnath H-O. Impact of correction factors in human brain lesion-behavior inference. Human Brain Mapping 2017; 38(3): 1692–701.

Sperber C, Wiesen D, Karnath H-O. An empirical evaluation of multivariate lesion behaviour mapping using support vector regression. Human Brain Mapping 2019; 40(5): 1381–90.

Takashima A, Konopka A, Meyer A, Hagoort P, Weber K. Speaking in the brain: The interaction between words and syntax in producing sentences. bioRxiv 2019: 696310.

Thompson CK, Bonakdarpour B, Fix SC, Blumenfeld HK, Parrish TB, Gitelman DR, et al. Neural Correlates of Verb Argument Structure Processing. Journal of Cognitive Neuroscience 2007; 19(11): 1753–67.

Thompson CK, Bonakdarpour B, Fix SF. Neural Mechanisms of Verb Argument Structure Processing in Agrammatic Aphasic and Healthy Age-matched Listeners. Journal of Cognitive Neuroscience 2010a; 22(9): 1993–2011.

Thompson CK, Cho S, Hsu C-J, Wieneke C, Rademaker A, Weitner BB, et al. Dissociations between fluency and agrammatism in primary progressive aphasia. Aphasiology 2012; 26(1): 20–43.

Thompson CK, Choy JJ, Holland A, Cole R. Sentactics®: Computer-automated treatment of underlying forms. Aphasiology 2010b; 24(10): 1242–66.

Thompson CK, den Ouden D-B. Neuroimaging and recovery of language in aphasia. Current Neurology and Neuroscience Reports 2008; 8(6): 475.

Thompson CK, Meltzer-Asscher A, Cho S, Lee J, Wieneke C, Weintraub S, et al. Syntactic and Morphosyntactic Processing in Stroke-Induced and Primary Progressive Aphasia. Behavioural Neurology 2013; 26(1-2).

Thothathiri M, Kimberg DY, Schwartz MF. The Neural Basis of Reversible Sentence Comprehension: Evidence from Voxel-based Lesion Symptom Mapping in Aphasia. Journal of Cognitive Neuroscience 2012; 24(1): 212–22.

Turkeltaub PE, Coslett HB, Thomas AL, Faseyitan O, Benson J, Norise C, et al. The right hemisphere is not unitary in its role in aphasia recovery. Cortex 2012; 48(9): 1179–86.

Tyler LK, Wright P, Randall B, Marslen-Wilson WD, Stamatakis EA. Reorganization of syntactic processing following left-hemisphere brain damage: does right-hemisphere activity preserve function? Brain 2010; 133(11): 3396–408.

Vigliocco G, Hartsuiker RJ. The interplay of meaning, sound, and syntax in sentence production. Psychological Bulletin 2002; 128(3): 442–72.

Weiller C, Isensee C, Rijntjes M, Huber W, Müller S, Bier D, et al. Recovery from wernicke’s aphasia: A positron emission tomographic study. Annals of Neurology 1995; 37(6): 723–32.

Welch BL. The Generalization of ‘Student’s’ Problem when Several Different Population Variances are Involved. Biometrika 1947; 34(1/2): 28–35.

Wilson SM, DeMarco AT, Henry ML, Gesierich B, Babiak M, Miller BL, et al. Variable disruption of a syntactic processing network in primary progressive aphasia. Brain 2016; 139(11): 2994–3006.

Wilson Stephen M, Galantucci S, Tartaglia Maria C, Rising K, Patterson Dianne K, Henry Maya L, et al. Syntactic Processing Depends on Dorsal Language Tracts. Neuron 2011; 72(2): 397–403.

Wilson SM, Henry ML, Besbris M, Ogar JM, Dronkers NF, Jarrold W, et al. Connected speech production in three variants of primary progressive aphasia. Brain 2010; 133(7): 2069–88.

Wilson SM, Saygın AP. Grammaticality Judgment in Aphasia: Deficits Are Not Specific to Syntactic Structures, Aphasic Syndromes, or Lesion Sites. Journal of Cognitive Neuroscience 2004; 16(2): 238–52.

Wu DH, Waller S, Chatterjee A. The Functional Neuroanatomy of Thematic Role and Locative Relational Knowledge. Journal of Cognitive Neuroscience 2007; 19(9): 1542–55.

Yourganov G, Fridriksson J, Rorden C, Gleichgerrcht E, Bonilha L. Multivariate Connectome-Based Symptom Mapping in Post-Stroke Patients: Networks Supporting Language and Speech. The Journal of Neuroscience 2016; 36(25): 6668–79.

Yourganov G, Fridriksson J, Stark B, Rorden C. Removal of artifacts from resting-state fMRI data in stroke. NeuroImage: Clinical 2018; 17: 297–305.

Zhang Y, Kimberg DY, Coslett HB, Schwartz MF, Wang Z. Multivariate lesion-symptom mapping using support vector regression. Human Brain Mapping 2014; 35(12): 5861–76.

Zingeser LB, Berndt RS. Retrieval of nouns and verbs in agrammatism and anomia. Brain and Language 1990; 39(1): 14–32.

